# Δ^9^-Tetrahydrocannabinol exposure shifts eosinophil and macrophage transcriptional programs towards an anti-inflammatory phenotype in helminth infection

**DOI:** 10.64898/2026.06.11.729983

**Authors:** Jennell Jennett, Martin Olmos, Ka Man Lam, Sarah Midou, Nicholas V. DiPatrizio, Meera G. Nair

## Abstract

Cannabis use is increasing globally, yet the immunological effects of Δ9-tetrahydrocannabinol (THC), the main intoxicating component of cannabis, remain incompletely understood. Given prior evidence that endocannabinoid signaling influences helminth immunity and type 2 inflammation, we investigated how sustained THC exposure alters immune responses to the helminth *Nippostrongylus brasiliensis* (Nb), which infects the lung and small intestine of mice. C57BL/6J mice were treated with THC (5 mg/kg/day) or vehicle for 14 days prior to helminth infection and assessed for parasite burden, innate immune cell and T cell responses, and transcriptional changes in lung eosinophils and macrophages. THC exposure did not significantly alter infection-associated weight loss or helminth burden; however, THC selectively restrained infection-induced circulating eosinophils and monocytes while increasing regulatory T cells. T cell activation assays showed reduced TNFα and IFNγ secretion in splenocytes from THC-treated infected mice. Bulk RNA sequencing showed that THC shifted lung eosinophils and CD11c⁺ lung macrophage-enriched cells from inflammatory, fibrotic, and costimulatory pathways toward stress and metabolic-adaptive transcriptional programs. Within the infected macrophage-enriched population, THC reduced CD80 expression while increasing MHC class II and antigen presentation-associated genes, suggesting a potential shift in macrophage-mediated T cell activation. Consistent with altered inflammatory and tissue remodeling-associated programs, immunofluorescent staining showed that THC mitigated infection-associated loss of lung collagen. Collectively, these findings indicate that THC reshapes the immune response to helminth infection by restraining innate and T cell effector responses while altering lung eosinophil and macrophage activation programs.

**Summary Sentence:** THC reshapes helminth-induced type 2 inflammation by restraining inflammatory leukocyte responses and reprogramming lung eosinophils and macrophages.

## Introduction

Cannabis is one of the most commonly used psychoactive substances in the United States. In 2022, an estimated 61.9 million people aged 12 years or older reported past year cannabis use, corresponding to 22.0% of the population in this age group. Cannabis use was highest among young adults aged 18 to 25, with 38.2% reporting past year use^1^. As cannabis use increases with expanding medical and recreational legalization, there is a growing need to understand how cannabis-derived compounds affect immune function. Δ^9^-tetrahydrocannabinol (THC) is the primary intoxicating component of cannabis and acts as a partial agonist at cannabinoid receptors^2^. Although THC is commonly studied for its neurological and behavioral effects, cannabinoid receptors and endocannabinoid pathways are also involved in immune regulation, raising important questions about how THC exposure alters inflammatory responses^34^.

The endocannabinoid system has emerged as an important regulator of host immunity during infection and inflammation. In helminth infection, both host- and helminth-derived endocannabinoids are generated and can influence host immune responses and parasite burden^5^. Human studies have also suggested a relationship between cannabis use and helminth infection. Among Aka foragers of the Congo Basin, cannabis use was highly prevalent and levels of urinary Δ^9^-tetrahydrocannabinol-11-oic acid (THC-COOH, the primary metabolite of THC) were negatively associated with parasite infection and reinfection^6^. However, this study evaluated whole cannabis exposure in a human population and could not determine whether THC directly altered parasite burden, host immunity, or both. Together, these findings suggest that cannabinoid pathways intersect with anti-helminth immunity, but the specific effects of THC on host immune responses remain unclear.

Infection with *Nippostrongylus brasiliensis*, a rodent hookworm, is a well-established murine model of helminth infection and type 2 inflammation, inducing eosinophilia, macrophage activation, Th2 cytokine responses, and lung tissue remodeling^7–9^. This model provides a controlled system to determine whether THC changes the magnitude or quality of immune responses during type 2 infection and inflammation. Prior work from our group showed that inhibition of peripheral cannabinoid receptor subtype 1 during *N. brasiliensis* infection increased airway eosinophilia and worsened pulmonary pathology, suggesting that cannabinoid signaling can restrain eosinophilic inflammation in this setting^10^. Considering that THC is a partial cannabinoid receptor agonist, these findings raise the possibility that THC exposure may alter endocannabinoid signaling at cannabinoid receptors and ensuing inflammatory responses during helminth infection-induced type 2 immunity.

THC has also been shown to regulate immune pathways relevant to inflammatory activation. In activated human T cells, THC suppresses proliferation, reduces IFNγ production, and alters Th1/Th2 cytokine balance^11^. In the myeloid compartment, THC has been reported to inhibit macrophage chemotaxis and alter macrophage differentiation through reactive oxygen species-dependent mechanisms^12–15^. These findings suggest that THC can regulate both adaptive and innate immune pathways. However, whether THC exposure alters the balance between immune activation, inflammatory restraint, and tissue remodeling during type 2 inflammation remains poorly understood.

Despite these observations, critical gaps remain. First, it is unclear whether THC alone is sufficient to alter parasite burden in a controlled helminth infection model, especially given human studies linking cannabis consumption to reduced helminth burden^6^. Second, it is unknown whether THC alters type 2 inflammation by reducing immune cell accumulation, changing inflammatory function, or modifying immune cell activation states. In addition, the consequences of THC-associated immune modulation for lung tissue repair following infection-induced injury remain unclear^9,16^. To address these gaps, we used *N. brasiliensis* infection as a controlled model of helminth infection and type 2 inflammation to evaluate how THC exposure affects parasite burden, innate and T cell immune responses, and overall outcomes in infection-induced lung tissue injury.

## Materials and Methods

### Mouse studies

All experiments were conducted in accordance with protocols approved by the Animal Care and Use Committee at the University of California, Riverside (AUP#48). Six-week-old male C57BL/6J mice were maintained in the vivarium under a 12-hour light, 12-hour dark cycle and provided ad libitum access to standard vivarium chow and water for the duration of the study.

Mice were treated daily with THC (5 mg/kg/day; generously donated by the National Institute on Drug Abuse Drug Supply Program) or vehicle for 14 days prior to infection. THC was dissolved in vehicle consisting of 7.5% DMSO, 7.5% Tween 80, and 85% sterile saline, and vehicle-treated mice received the corresponding vehicle control. On day 14, mice were briefly anesthetized with isoflurane and infected subcutaneously with 550 *Nippostrongylus brasiliensis* L3 larvae, as previously described for murine lung-stage helminth infection models ^7,9^. THC and vehicle injections were paused from the day of infection until 4 days post infection to minimize handling during the acute phase of infection. Daily THC or vehicle treatment resumed on day 18 and continued until the study endpoint on day 22. Fecal samples were collected at day 7 post infection, and egg burden was quantified as eggs per gram of feces. At day 8 post infection, mice were euthanized and blood, bronchoalveolar lavage fluid, lungs, spleens, and small intestines were collected for downstream analyses. Adult worm burden was quantified from the small intestine. THC was obtained through a collaboration with the laboratory of Dr. Nicholas DiPatrizio at the University of California, Riverside, under applicable institutional approvals and federal controlled substance regulations. THC was stored, prepared, and administered by authorized personnel in accordance with DEA controlled substance requirements and UCR institutional procedures. All animal procedures were approved by the University of California, Riverside Institutional Animal Care and Use Committee under protocol 48.

### BALF, lung, and blood leukocyte isolation and flow cytometry

Bronchoalveolar lavage fluid (BALF) was collected by three lavages of 0.7 mL sterile 1X PBS. Whole lungs were dissociated using the mouse Lung Dissociation Kit (Miltenyi Biotec, Cat. 130-095-927) with the gentleMACS Octo Dissociator with Heaters according to the manufacturer’s protocol. Blood was collected into 4% sodium citrate and diluted with wash media prior to density gradient separation with Histopaque-1077 (Sigma-Aldrich, Cat. 10771). Following centrifugation at 400 × g for 20 minutes at room temperature with the brake off, the leukocyte fraction was collected, washed, and resuspended for flow cytometry. For BALF, lung, and blood leukocyte suspensions, viable cells were counted on the DeNovix CellDrop, followed by standard staining for flow cytometry included CD16/32 (BioLegend, 5 µg/mL) and rat IgG (Millipore Sigma, 10 µg/mL), followed by staining with surface antibody cocktail (0.5 µg/mL/antibody; Supplementary Table S1, S2). Data were analyzed using NovoExpress and FlowJo flow cytometry analysis software.

### Ex-vivo splenocyte stimulation

Spleens were collected into wash media and mechanically dissociated through a cell strainer to generate single-cell suspensions. Cells were centrifuged at 400 × g for 5 minutes at 4°C, followed by red blood cell lysis using RBC Lysis Buffer (BioLegend, Cat. 420302) for 5 minutes. Splenocytes were washed, counted, and resuspended in complete T cell culture media consisting of DMEM supplemented with 10% FBS, 1% penicillin-streptomycin, HEPES, and β- mercaptoethanol. Splenocytes were plated at 5 × 10⁶ cells per well and stimulated with anti-CD3 (clone 17A2, eBioscience, Cat. 16-0032-82) and anti-CD28 (clone 37.51, eBioscience, Cat. 16-0281-82) antibodies at 1 µg/mL each, with recombinant mouse IL-2 at 15 ng/mL (Gibco/PeproTech, Cat. 21212100UG), for 72 hours at 37°C. Following incubation, culture supernatants were collected and stored at -20°C until cytokine analysis. IFNγ was measured using the LEGENDplex™ Mouse Th1/Th2 Panel 8-plex with V-bottom Plate (BioLegend, Cat. 741054). TNFα was measured using the Mouse TNF-alpha Quantikine ELISA Kit (R&D Systems, Cat. MTA00B).

### Lung macrophage and eosinophil isolation for bulk RNA sequencing

Whole lung single cells were resuspended in cold MACS buffer and filtered through 35 µm flow cytometry tube strainers to remove remaining aggregates followed by magnetic bead purification. Cells were incubated with biotinylated anti-mouse CD11c antibody (BioLegend, clone N418, Cat. 117304) at 1:100 in MACS buffer for 10 minutes at 4°C, followed by incubation with Streptavidin MicroBeads (Miltenyi Biotec, Cat. 130-048-102). Labeled cells were passed over MS columns, and retained CD11c+ cells were eluted for downstream RNA isolation. The CD11c-negative flow-through was retained for eosinophil enrichment with Anti-Siglec-F MicroBeads (Miltenyi Biotec, Cat. 130-118-513) and MS column separation. Enriched CD11c+ macrophage and Siglec-F+ eosinophil fractions were counted, pelleted, and lysed in Buffer RLT. RNA was isolated using the RNeasy Mini Kit (QIAGEN, Cat. 74106) according to the manufacturer’s protocol. Briefly, cell lysates were mixed with 70% ethanol and applied to RNeasy spin columns. Columns were washed with RW1 and RPE buffers, followed by an additional 80% ethanol wash. RNA was eluted in RNase-free water prewarmed to 60°C. RNA concentration and purity were assessed by spectrophotometry, and samples were submitted to Novogene Corporation, Inc. for library preparation and bulk RNA sequencing. Raw sequencing reads were assessed for quality using FastQC ^52^and summarized with MultiQC^53^. Adapter sequences and low-quality bases were trimmed using Trimmomatic^54^. Trimmed reads were aligned to the mouse reference genome using STAR aligner^55^, and gene-level read counts were generated using FeatureCounts^56^. Downstream analyses were performed in R. Differential gene expression analysis was conducted using DESeq2^57^. Genes with an adjusted p value, using Benjamini-Hochberg correction, ≤ 0.05 and an absolute log2 fold change > 1 were considered differentially expressed unless otherwise indicated. For targeted antigen presentation and costimulation-associated genes, an adjusted p value ≤ 0.05 and absolute log2 fold change > 0.25 were used. Variance-stabilized expression values generated by DESeq2 were used for heatmap visualization. Gene set enrichment analysis was performed using Hallmark gene sets^58,59^.

### Lung immunofluorescent staining and imaging

Lungs were inflated with 4% paraformaldehyde in PBS, collected, and fixed overnight in 10% neutral buffered formalin at 4°C. Tissues were then transferred to 70% ethanol and submitted to the Translational Pathology Core Laboratory at UCLA for paraffin embedding and sectioning. Lung sections were cut at 5 µm. For immunofluorescent staining, FFPE lung sections were dewaxed and rehydrated with standard xylene, ethanol, and distilled water washes, followed by antigen retrieval by boiling slides in 1× citrate antigen retrieval buffer, pH 6.0, prepared from 10× Citrate Buffer pH 6.0 (Abcam, Cat. ab64214) for 20 minutes. Sections were encircled with a hydrophobic barrier pen, permeabilized in 0.4% Triton X-100 in PBS, and blocked in 5% goat serum and 2.5% BSA in PBST. Avidin and biotin blocking were performed using the Avidin/Biotin Blocking Kit (Vector Laboratories, Cat. SP-2001) according to the manufacturer’s instructions. Sections were incubated overnight at 4°C with primary antibodies against Collagen VI and GSL I (Supplementary Table S3), diluted in PBST supplemented with 2.5% BSA. Sections were washed twice in PBST, incubated with secondary antibodies for 2 hours at room temperature, washed twice in PBST, and rinsed briefly in distilled water. Following secondary antibody incubation, sections were treated with ReadyProbes™ Tissue Autofluorescence Quenching Kit according to the manufacturer’s instructions (Invitrogen, Cat. No. R37630). Sections were mounted with VECTASHIELD Vibrance Antifade Mounting Medium with DAPI (Vector Laboratories, Cat. H-1800-2). Immunofluorescent lung sections were imaged at 20× magnification using a Keyence microscope. Collagen VI staining was imaged in the Cy5 channel and GSL I was imaged in the Cy3 channel. Collagen VI signal was quantified using Fiji/ImageJ. Images were converted to 16-bit grayscale, background was subtracted, and Collagen VI integrated density was measured across the full field of view for each image. Data are reported as Collagen VI integrated density per field.

## Results

### THC exposure does not alter parasite burden following *Nippostrongylus brasiliensis* infection but affects circulating leukocyte responses

Cannabis use has been associated with altered helminth burden in human population^6^, raising the possibility that cannabinoid exposure may influence host parasite responses. To determine whether THC exposure altered helminth burden and infection-associated immune responses, we employed an established murine model of helminth infection with *Nippostrongylus brasiliensis* (Nb), a rodent hookworm that infects the lung and small intestine with an acute infection cycle of 7 to 10 days^7,17,18^. Mice were treated with THC or vehicle for two weeks prior to Nb infection, followed by assessment of infection-associated weight change, fecal egg burden, and intestinal worm burden (Fig. 1A). Following Nb infection, vehicle and THC-treated mice showed acute weight loss between days 2 and 4 post-infection, consistent with the timing of Nb-associated lung migration and tissue injury (Fig. 1B). However, THC treatment did not significantly alter fecal egg burden or adult worm burden compared with vehicle-treated infected controls (Fig. 1C, D).

**Figure 1.**
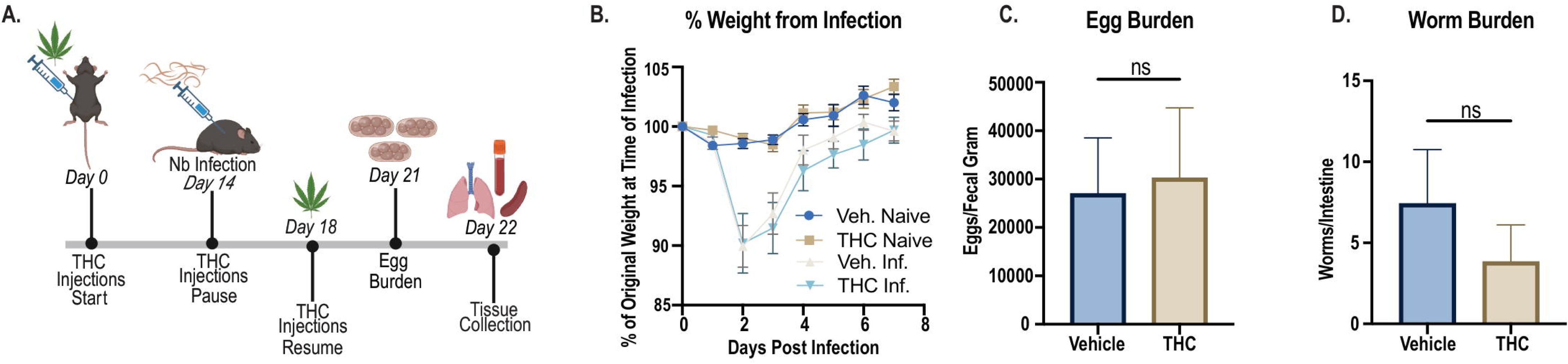
THC exposure does not alter infection-induced weight loss or parasite burden following Nippostrongylus brasiliensis infection. Vehicle and THC-treated mice were left naïve or infected with Nb and assessed for infection-associated weight change and parasite burden at day 7 post-infection. (A) Experimental timeline. (B) Body weight change shown as percent of original body weight at the time of infection. (C) Fecal egg burden in vehicle and THC-treated infected mice. (D) Adult worm burden in the small intestine. Data are presented as mean ± SEM of n= 7 per group and representative of two independent experiments. Statistical analysis was performed using an unpaired t test for egg burden and worm burden. ns, not significant.

We next examined whether THC altered circulating immune cell responses during Nb infection. Whole blood flow cytometry was performed to quantify monocytes, eosinophils and CD4^+^ T cells (Fig. S1). Nb infection increased circulating eosinophil frequencies in vehicle-treated mice, whereas this infection-induced eosinophil response was reduced in THC-treated animals (Fig. 2A). Nb infection also increased circulating monocyte frequencies in vehicle-treated mice, and THC-treated infected mice had reduced circulating monocyte frequencies compared with vehicle-treated infected controls (Fig. 2B). In contrast, CD4^+^CD25^+^ regulatory T cell frequencies were increased in THC-treated infected mice (Fig. 2C). Total CD4+ T cell frequencies in the blood were not significantly altered by THC treatment (Fig. S1C), indicating that THC did not broadly impact the proportion of circulating CD4+ T cells.

**Figure 2.**
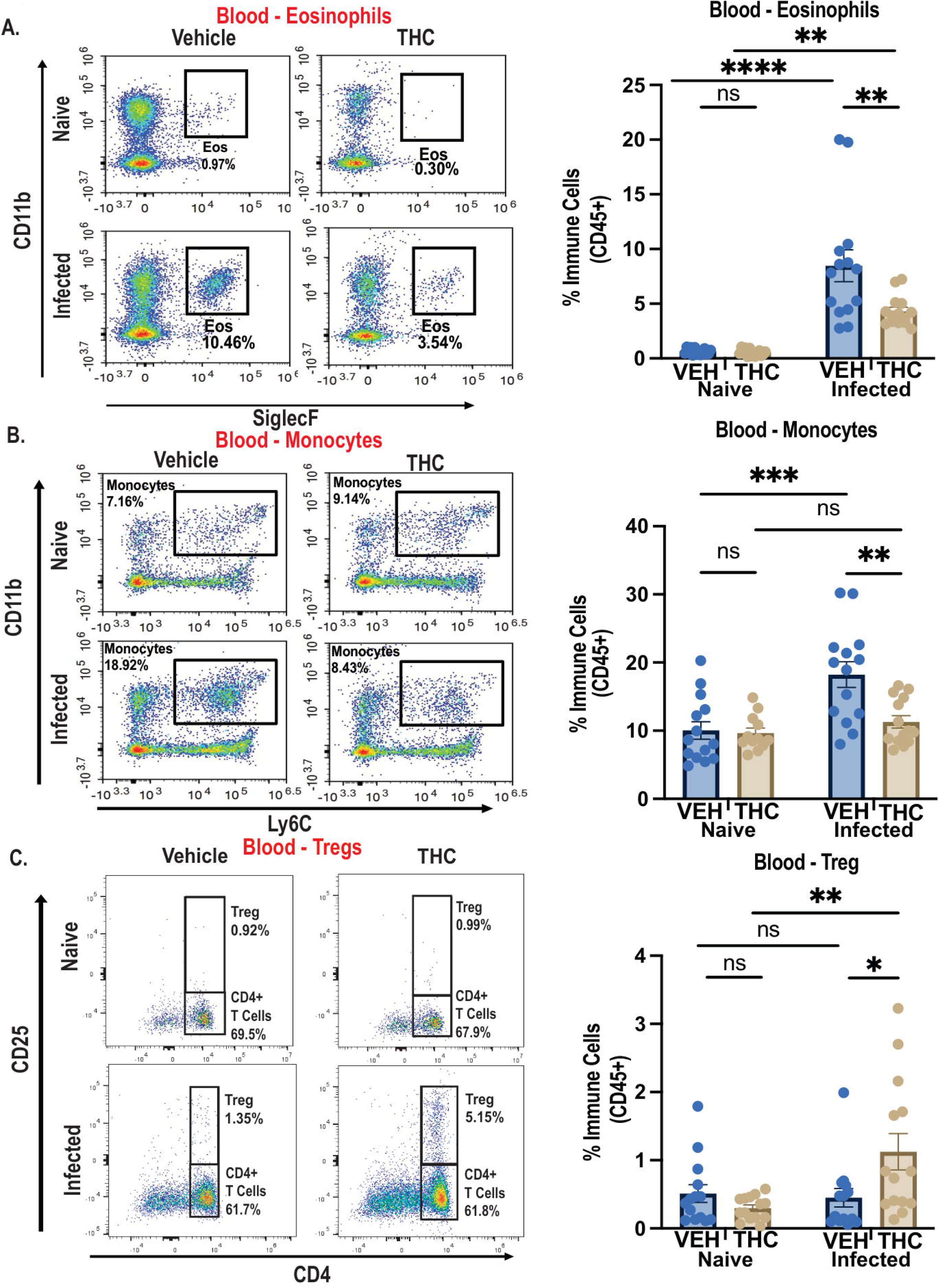
THC exposure reduces infection-induced circulating eosinophilia and alters circulating leukocyte responses. Whole blood leukocyte responses were assessed in vehicle and THC-treated naïve and Nb-infected mice. (A-C) Representative flow cytometry plots and frequency of eosinophils (A), monocytes (B) and CD4⁺CD25⁺ regulatory T cells (C) within CD45⁺ cells. Data are presented as mean ± SEM and pooled from 2 independent experiments with n = 4 to 7 mice per group. Statistical analysis was performed using two-way ANOVA with Tukey’s multiple comparisons test. *p < 0.05; **p < 0.01; ***p < 0.001; ****p < 0.0001; ns, not significant.

Together, these data indicate that THC exposure does not significantly alter parasite burden or infection-associated weight loss following Nb infection but does alter systemic immune responses. Specifically, THC limited infection-induced circulating eosinophilia and monocyte responses while increasing regulatory T cell frequencies. Overall, these findings support a model in which THC does not broadly suppress anti-helminth immunity but instead shifts the balance of systemic inflammatory and regulatory immune responses during type 2 inflammation.

### THC and helminth infection induce broad transcriptional changes in lung CD11c+ enriched macrophages and eosinophils

Although THC reduced circulating eosinophils and monocytes (Fig. S2), lung eosinophil and myeloid cell numbers in naïve mice or following Nb infection were not significantly changed with THC exposure (Fig. S3). We therefore investigated whether THC altered the gene expression profiles of these innate immune cells within the lungs of naïve and Nb-infected mice that had been exposed to vehicle or THC by magnetic column purification then bulk RNA sequencing (Fig. 3A). Lung single cell suspensions were processed for column purification to isolate CD11c^+^ alveolar macrophages as the dominant population, as well as dendritic cells (DC) (86% purity). In the infected mice groups, which had sufficient eosinophil numbers, the flow through was next used for SiglecF column purification to isolate eosinophils (87% purity).

**Figure 3.**
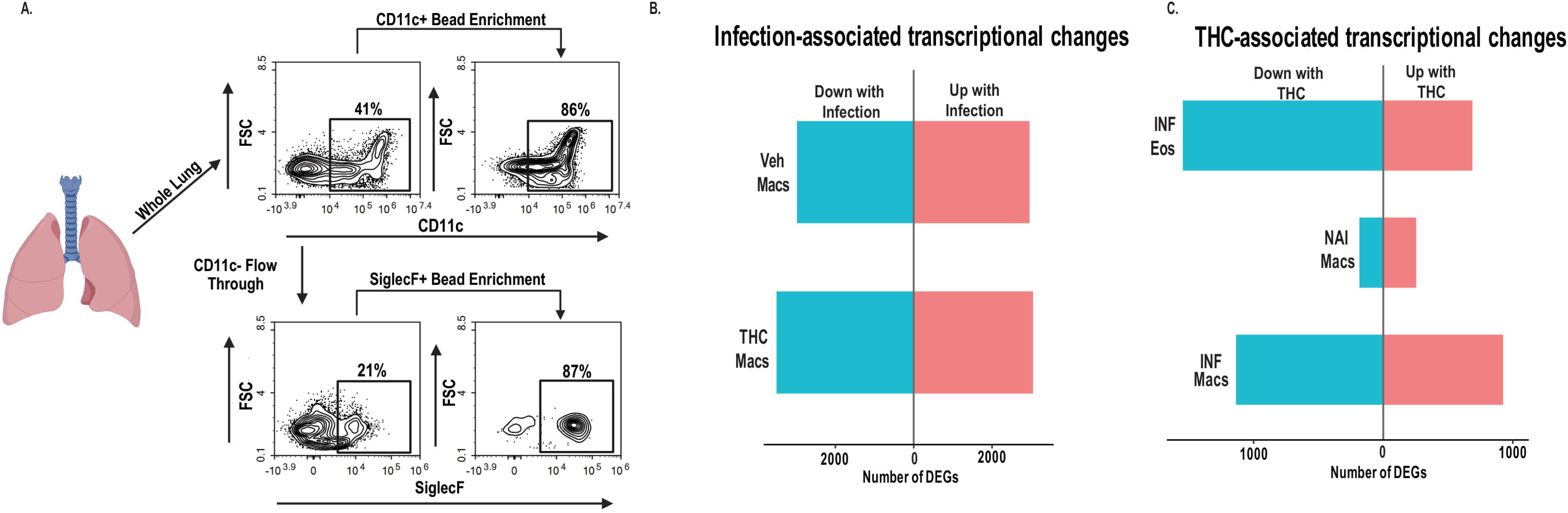
THC exposure and helminth infection induce broad transcriptional changes in lung macrophage enriched CD11c^+^ cells and eosinophils. Vehicle and THC-treated mice were left naïve or infected with Nb for 7 days. (A) Magnetic cell sorting strategy used to isolate CD11c⁺ lung macrophage-enriched cells and CD11c⁻Siglec-F⁺ eosinophils from whole lung homogenates for bulk RNA sequencing. (B) Differentially expressed gene counts comparing naïve and infected CD11c⁺ lung macrophage-enriched cells from vehicle and THC-treated mice. (C) Differentially expressed gene counts comparing vehicle and THC-treated infected eosinophils, naïve CD11c⁺ lung macrophage-enriched cells, and infected CD11c⁺ lung macrophage-enriched cells. Differentially expressed genes were identified by DESeq2 using an adjusted p value < 0.05 and absolute log2 fold change > 1. Data are representative of 1 experiment with 7 mice per group pooled into 4 RNA sequencing libraries per group.

Following bulk RNA sequencing and standard quality control filtering, we assessed the overall transcriptional impact of infection and THC exposure, by comparing the number of differentially expressed genes across RNA sequencing datasets (n=4 per group). In CD11c^+^ lung macrophage-enriched cells, Nb infection induced the greatest transcriptional changes in both vehicle and THC-treated mice; vehicle-treated macrophage-enriched cells had 2,956 genes upregulated and 2,980 genes downregulated with infection, compared to THC-treated macrophage-enriched cells (3,045 genes up versus 3,502 genes down with infection) (Fig. 3B). While THC did not substantially affect macrophage gene expression from naïve mice (254 genes up versus 184 genes down with THC), there were more genes that were changed in the infected mouse groups (Fig. 3C). Eosinophils had more downregulated (1546) than upregulated genes (688), followed by macrophages with 1,136 genes downregulated versus 925 genes upregulated with THC exposure in the infected groups. Together, these data indicate that THC has a broad transcriptional impact on macrophages and eosinophils during infection with a greater effect on suppressing gene expression.

Given prior evidence that THC can alter macrophage inflammatory potential^12–14^, we examined the genes and functional pathways that were changed in macrophages from naïve mice upon THC exposure (Fig. 4, Table 1). Transcripts reduced in THC-treated naïve cells included chemokine and inflammatory signaling-associated genes such as *Cxcl1, Cxcl2, Il2rg, Cish,* and *Scimp*, as well as heat shock-associated genes including *Hsph1* and *Hspa8*^31–36^. In contrast, genes increased with THC included *Cirbp, Cd300lf, Adap2, Rgcc, Pex11a, Cerk,* and *Pla2g6*, consistent with altered stress adaptation, inhibitory signaling, and lipid handling^21,26,37^.

**Figure 4.**
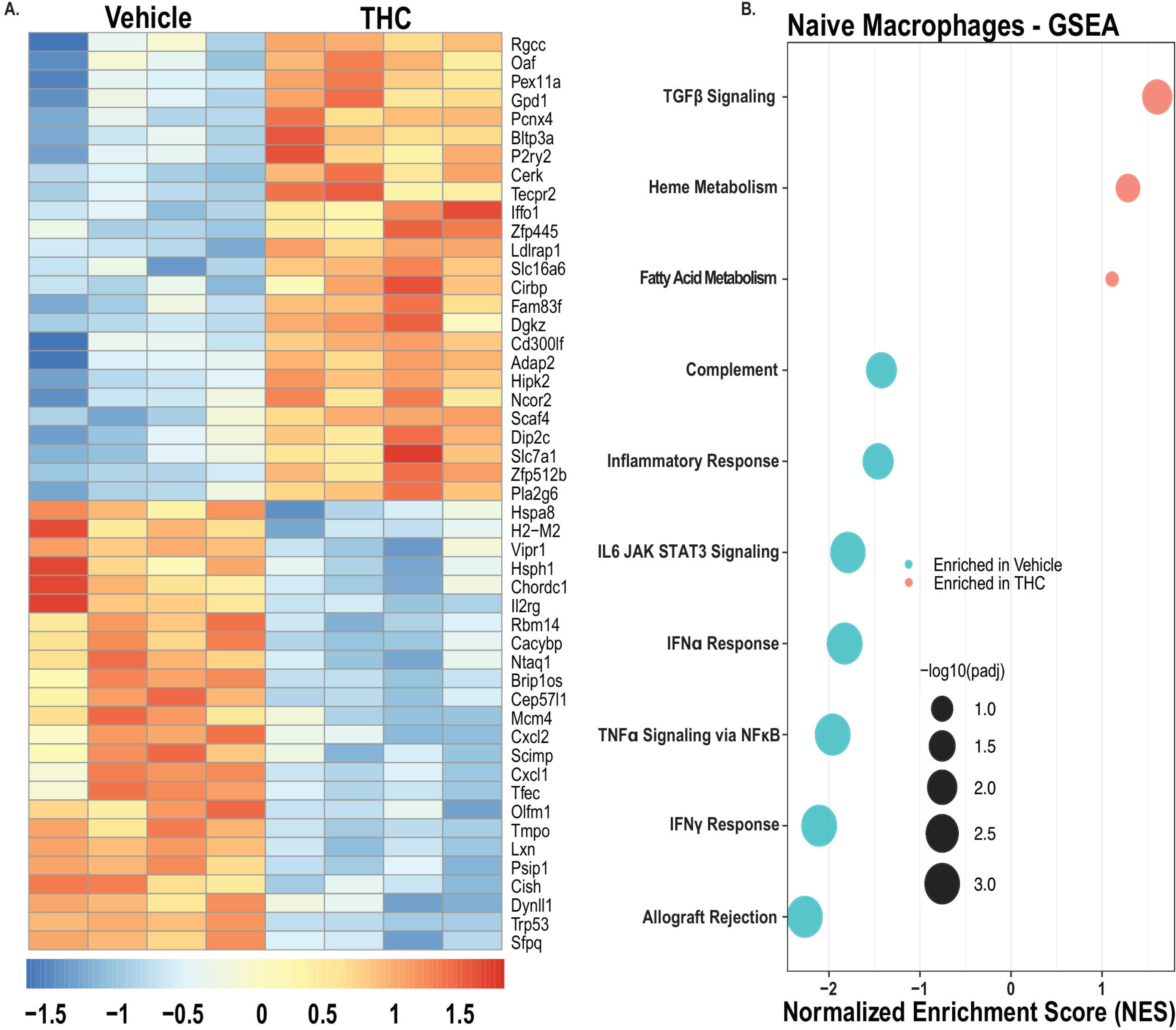
THC reduces basal inflammatory programming in CD11c⁺ lung macrophage enriched cells. CD11c⁺ lung macrophage-enriched cells from naïve vehicle and THC-treated mice were analyzed by bulk RNA sequencing. (A) Heatmap of the top 50 differentially expressed genes. (B) Hallmark gene set enrichment analysis. Genes shown in (A) met criteria of absolute log2 fold change > 1 and adjusted p value < 0.05 by DESeq2. GSEA pathways shown in (B) met adjusted p value < 0.05. Data are representative of 1 experiment with 7 mice per group pooled into 4 RNA sequencing libraries per group.

**Table. 1.** Naive lung CD11c+ enriched macrophage differentially expressed genes.

| Category | Down in THC | Up in THC | Interpretation |
| --- | --- | --- | --- |
| **Stress response** | *Hsph1, Hspa8, Chordc1* | *Cirbp* | THC reduces classical heat-shock stress, enhances stress-buffering |
| **Chemokines** | *Cxcl1, Cxcl2* | — | Vehicle macs show higher basal chemokine production |
| **Cytokine signaling & activation** | *Il2rg, Cish, Tfec, Scimp* | *Cd300lf, Adap2* (inhibitory) | THC upregulates inhibitory receptors, lowering activation potential |
| **Cell cycle / proliferation** | *Mcm4, Tmpo, Psip1* | *Rgcc, Pex11a, Gpd1, Slc16a6* | THC shifts toward metabolic/FAO genes rather than proliferation |
| **Lipid turnover** | — | *Cerk, Pla2g6* | THC enhances lipid-handling without inflammatory activation |

Gene set enrichment analysis supported this basal shift in CD11c+ lung macrophage-enriched cells. THC-treated naïve cells showed enrichment of pathways associated with TGFβ signaling, heme metabolism, and fatty acid metabolism. In contrast, vehicle-treated cells were enriched for inflammatory response, TNFα signaling via NFκB, IL-6 JAK STAT3 signaling, interferon responses, complement, and allograft rejection pathways (Fig. 4B).

We next examined whether THC altered CD11c+ lung macrophage-enriched cell responses during Nb infection. Differential gene expression analysis showed that THC altered the transcriptional response of infected CD11c+ lung macrophage-enriched cells (Fig. 5A, Table 2). THC reduced expression of genes associated with pathogen sensing, chemotaxis, motility, and remodeling. Reduced transcripts included *Tlr4, Fpr1,* and *P2ry13*, as well as heat shock-associated genes such as *Hsph1, Hsp90aa1,* and *Hspa8*^31,38–41^. THC also reduced genes linked to cytoskeletal remodeling and fibrotic programs, including *Myo1f, Bin2, Spice1,* and *P4ha1*^42–45^. Conversely, THC increased expression of genes associated with stress adaptation, regulatory signaling, and metabolic remodeling, including *Cirbp, Rbm3, Foxred2, Eci2, Pik3ip1, Lpxn, Susd3, Retnlg,* and *Pdcd4*^21,22,46–48^.

**Figure 5.**
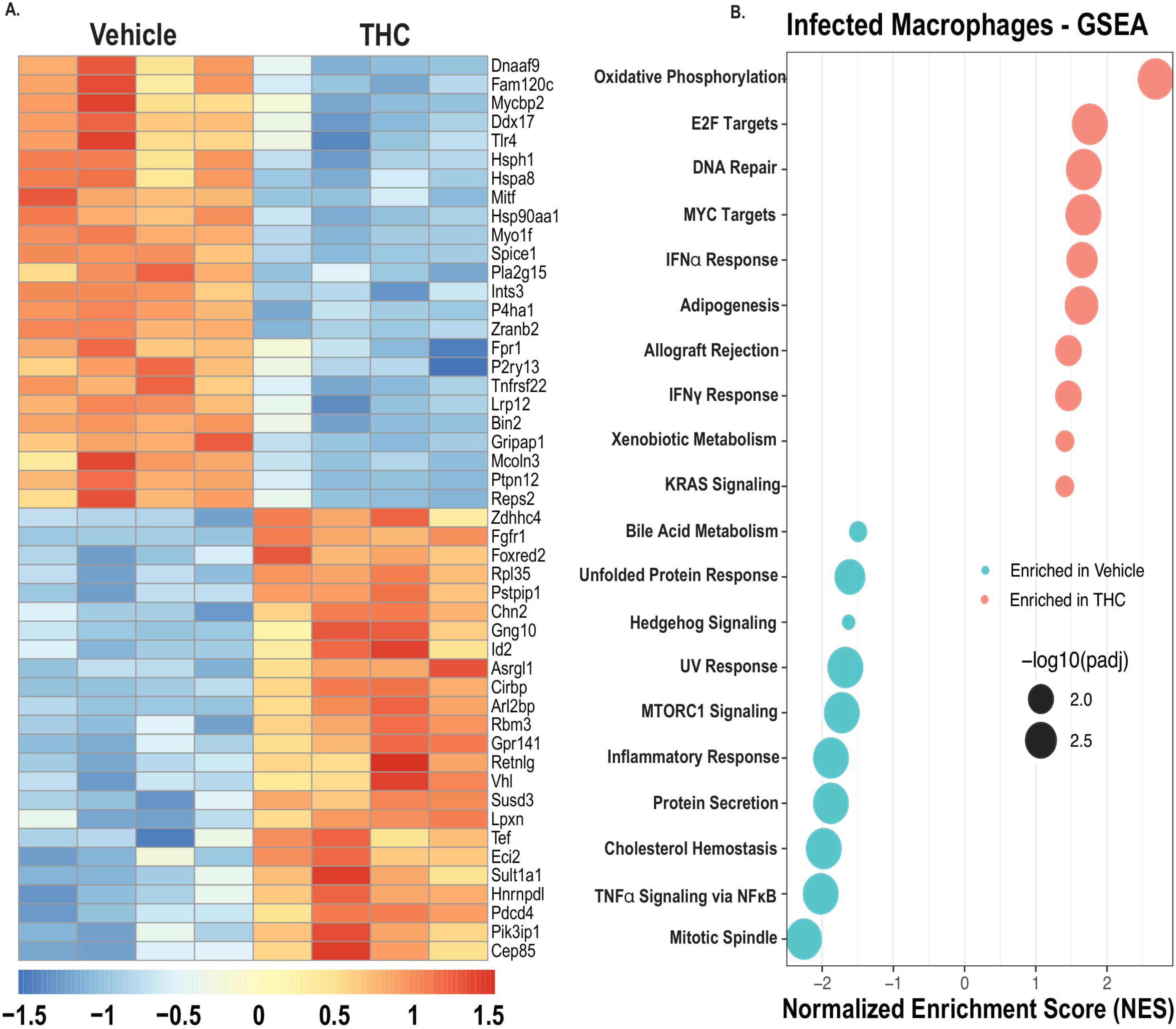
THC dampens inflammatory and remodeling macrophage programs during Nb induced inflammation. CD11c⁺ lung macrophage-enriched cells from vehicle and THC-treated Nb-infected mice were analyzed by bulk RNA sequencing. (A) Heatmap of the top 50 differentially expressed genes. (B) Hallmark gene set enrichment analysis. Genes shown in (A) met criteria of absolute log2 fold change > 1 and adjusted p value < 0.05 by DESeq2. GSEA pathways shown in (B) met adjusted p value < 0.05. Data are representative of 1 experiment with 7 mice per group pooled into 4 RNA sequencing libraries per group.

**Table 2.** Infected lung CD11c+ enriched macrophage differentially expressed genes.

| Category | Down in THC | Up in THC | Interpretation |
| --- | --- | --- | --- |
| **Pathogen sensing / chemotaxis** | *Tlr4, Fpr1, P2ry13* | — | THC reduces PRR/chemotaxis signaling |
| **Stress response** | *Hsph1, Hsp90aa1, Hspa8* (heat-shock proteins) | *Cirbp, Rbm3, Foxred2* (cold-shock/stress-buffering) | THC shifts from inflammatory stress responses to protective stress-buffering |
| **Metabolism** | *Pla2g15* (lipid metabolism for inflammation) | *Eci2* (FAO) | THC redirects metabolism away from inflammatory lipid use toward FAO |
| **Cytoskeletal/adhesion** | *Myo1f, Bin2, Spice1* | — | Reduced motility/adhesion signatures |
| **Fibrosis/ECM remodeling** | *P4ha1* | — | Lower fibrosis/ECM remodeling potential |
| **Immune regulation** | — | *Pik3ip1, Lpxn, Susd3, Retnlg, Pdcd4* | THC increases inhibitory/checkpoint signals |

Pathway analysis confirmed that THC altered macrophage-enriched transcriptional responses during infection. In vehicle-treated macrophages from infected mice, pathways associated with inflammatory response, TNFα signaling via NFκB, mTORC1 signaling, protein secretion, unfolded protein response, cholesterol homeostasis, and mitotic spindle were enriched. On the other hand, THC exposure promoted the enrichment of genes associate with oxidative phosphorylation, E2F targets, DNA repair, Myc targets, interferon responses, adipogenesis, and xenobiotic metabolism pathways (Fig. 5B). Together, these macrophage-enriched datasets from the lungs of naïve and Nb-infected mice indicate that THC reduces inflammatory, chemokine-associated, and remodeling programs, especially with infection, and instead promotes transcriptional features associated with metabolic adaptation, stress regulation, and immune restraint.

Although THC reduced circulating eosinophilia, lung eosinophil accumulation following Nb infection was not significantly reduced with THC exposure (Fig. S2C). We therefore asked whether THC altered the transcriptional state of eosinophils that reached the lung during infection. Differential gene expression analysis revealed that THC altered the transcriptional profile of infected lung eosinophils (Fig. 6A, Table 3). Among the top 50 differentially expressed genes, THC-treated eosinophils showed increased expression of transcripts associated with stress adaptation and cellular buffering, including *Ucp2, Cirbp, Rbm3,* and *Gadd45a*^19–23^. THC-treated eosinophils also showed increased expression of genes associated with lipid mediator regulation and inhibitory receptor signaling, including *Pla2g7, Cyp4f18, Cd300lf,* and *Siglecf*^24–28^. In contrast, transcripts associated with adhesion, migration, and inflammatory stress, including *Macf1, Syne2, Afap1l2, Neat1,* and *Hsph1*, were reduced in THC-treated eosinophils^29–31^.

**Figure 6.**
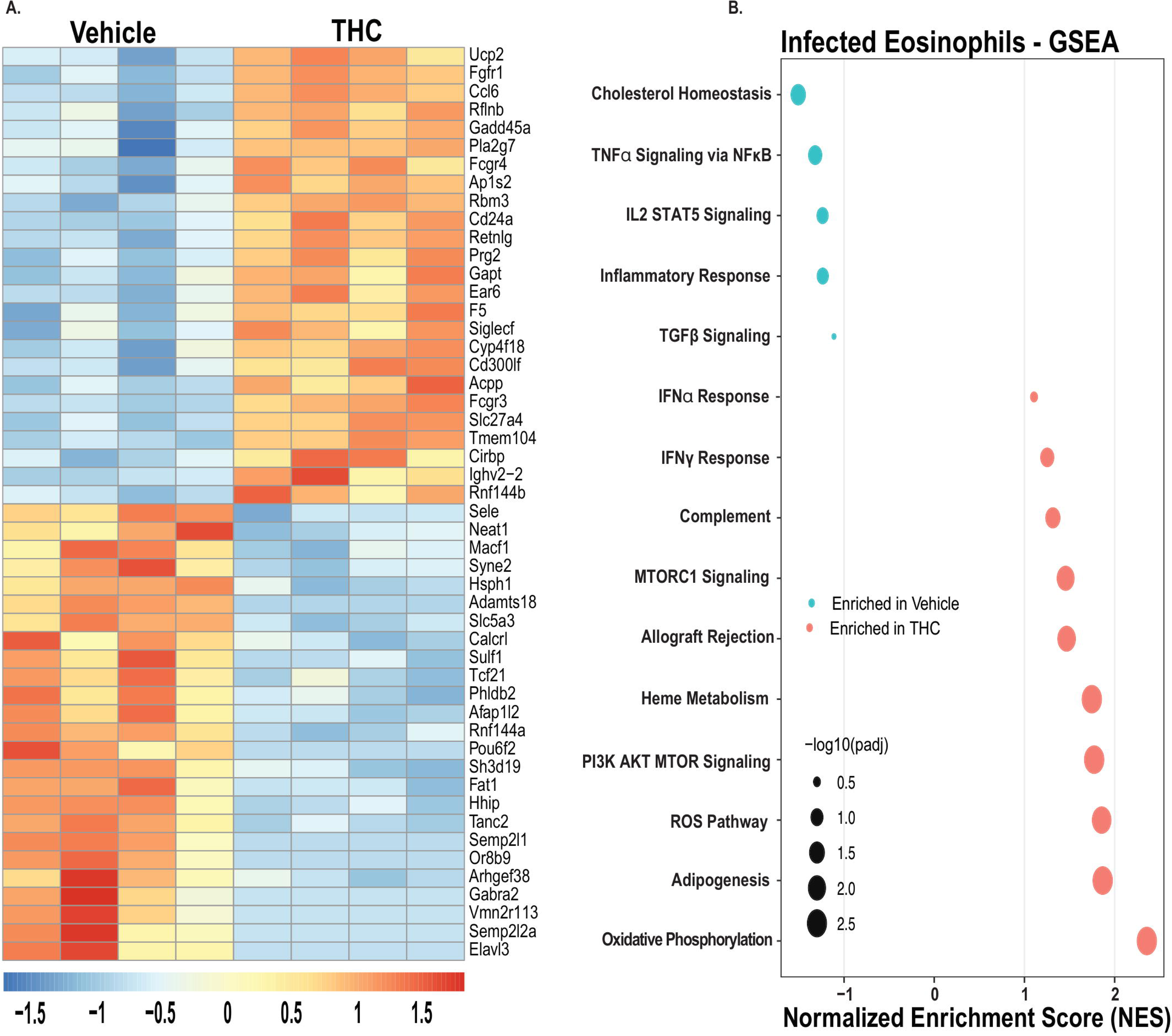
THC shifts infected lung eosinophils from inflammatory NFκB/STAT signaling to mTOR signaling and oxidative phosphorylation. CD11c⁻Siglec-F⁺ lung eosinophils from vehicle and THC-treated Nb-infected mice were analyzed by bulk RNA sequencing. (A) Heatmap of the top 50 differentially expressed genes. (B) Hallmark gene set enrichment analysis. Genes shown in (A) met criteria of absolute log2 fold change > 1 and adjusted p value < 0.05 by DESeq2. GSEA pathways shown in (B) met adjusted p value < 0.05. Data are representative of 1 experiment with 7 mice per group pooled into 4 RNA sequencing libraries per group.

**Table 3.** CD11c-SiglecF+ enriched lung eosinophil differentially expressed genes.

| Category | Down in THC | Up in THC | Interpretation |
| --- | --- | --- | --- |
| **Stress response / antioxidant** | — | *Ucp2, Cirbp, Rbm3, Gadd45a* | Enhanced buffering of oxidative and cellular stress |
| **Lipid mediator metabolism** | — | *Pla2g7, Cyp4f18* | Reduced PAF/LTB4 signaling capacity (neutrophil recruitment) |
| **Inhibitory receptors** | — | *Cd300lf, Siglec f* | Higher activation threshold |
| **Cytoskeleton / adhesion** | *Macf1, Syne2, Afap1l2* | — | Lower migratory and adhesion potential |
| **Stress / inflammatory regulators** | *Neat1, Hsph1* | — | Reduced stress-induced inflammatory signaling |

Gene set enrichment analysis further supported a THC-associated shift in eosinophil activation state. Pathways associated with oxidative phosphorylation, adipogenesis, reactive oxygen species, PI3K AKT mTOR signaling, and heme metabolism were enriched in THC-treated eosinophils (Fig. 6B). In contrast, inflammatory response, TNFα signaling through NFκB, IL-2 STAT5 signaling, TGFβ signaling, and cholesterol homeostasis pathways were enriched in vehicle-treated eosinophils. Together, these data indicate that THC does not primarily limit eosinophil accumulation in the lung during Nb infection but instead alters the transcriptional state of lung eosinophils that enter the tissue. Here, THC may restrain eosinophil inflammatory potential at two levels: by reducing the circulating eosinophil response and by shifting infected lung eosinophils toward a less inflammatory, more antioxidant-associated transcriptional state after tissue entry.

### THC alters antigen presentation and costimulatory gene expression while limiting inflammatory T cell activation

Given that THC altered circulating leukocyte responses and triggered anti-inflammatory transcriptional programs in CD11c+ lung macrophages, we hypothesized that inflammatory T cell responses may be changed. Flow cytometric analysis of BALF showed that THC-treated infected mice had increased CD4^+^ T cell frequencies compared with vehicle-treated infected mice (Fig. 7A, B). To determine whether this increase corresponded with enhanced inflammatory T cell function, splenocytes from vehicle and THC-treated mice were stimulated ex vivo with anti-CD3/CD28 and IL-2 for 72 hours, and cytokine secretion was measured in culture supernatants (Fig. 7C). Splenocytes from THC-treated infected mice generated significantly less IFNγ and TNFα following stimulation compared with splenocytes from vehicle-treated infected controls (Fig. 7D, E). These data indicate that THC increased CD4^+^ T cell accumulation in the BALF; however, its systemic impact was immunosuppressive, characterized by decreased blood Tregs and reduced inflammatory cytokine production by splenic T cells.

**Figure 7.**
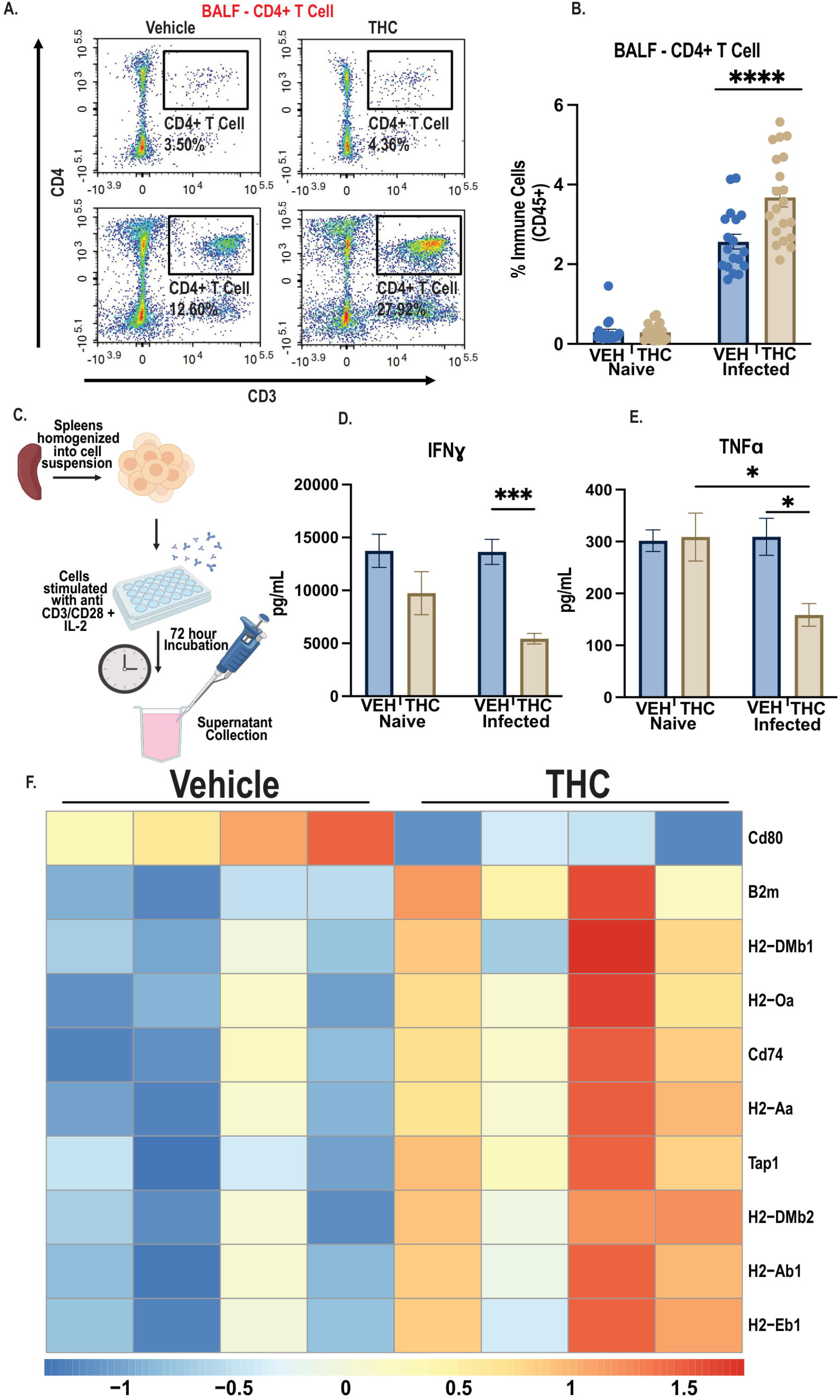
THC limits inflammatory T cell activation while preserving antigen presentation programs. T cell accumulation, cytokine production, and antigen presentation-associated gene expression were assessed in vehicle and THC-treated mice. (A and B) Representative BALF flow cytometry plots (A) and frequency of CD4⁺ T cells among CD45⁺ cells (B) from naïve and Nb-infected mice. (C) Schematic of ex vivo splenocyte stimulation. (D and E) IFNγ and TNFα concentrations following ex vivo splenocyte stimulation. (F) Targeted heatmap of MHC class II, antigen presentation, and costimulation-associated genes in CD11c⁺ lung macrophage-enriched cells from vehicle and THC-treated infected mice. Data are presented as mean ± SEM. BALF flow cytometry and cytokine data are pooled from 2 independent experiments with n = 4 to 7 mice per group. RNA sequencing data are representative of 1 experiment with 7 mice per group pooled into 4 RNA sequencing libraries per group. Statistical analysis was performed using two-way ANOVA with Tukey’s multiple comparisons test. Genes shown in (F) met criteria of absolute log2 fold change > 0.25 and adjusted p value < 0.05 by DESeq2. *p < 0.05; ***p < 0.001; ****p < 0.0001.

Given that antigen-presenting cells regulate T cell activation through MHC-dependent antigen presentation and CD80/CD86-mediated costimulatory signaling^49^, we next examined antigen presentation and co-stimulation-associated genes in CD11c+ lung macrophages. Under naïve conditions, there were no significant differences in antigen presentation or co-stimulation-associated genes between vehicle and THC-treated mice. During infection, however, THC-treated CD11c+ lung macrophages maintained or increased expression of multiple MHC class II and antigen presentation-associated genes, including *H2-Aa, H2-Ab1, H2-Eb1, Cd74, H2-DMb1, H2-DMb2, H2-Oa, B2m,* and *Tap1*^50^(Fig. 7F). In contrast, *Cd80* expression was reduced in THC- treated infected cells, while *Cd86* was assessed but not significantly changed. Together, these data indicate that THC-treated mice have increased airway CD4+ T cell frequencies but reduced inflammatory cytokine output following stimulation. This pattern suggests that THC does not block antigen presentation or fully eliminate costimulatory potential, but may partially dampen costimulatory signaling in a manner that limits inflammatory T cell activation^12^.

### THC treatment is associated with retained lung Collagen VI following Nb infection

Based on the transcriptomic profiling results, showing that THC altered inflammatory and remodeling-associated transcriptional programs in lung macrophage-enriched cells, we next assessed whether THC was associated with tissue-level changes in the lung. We chose to investigate collagen deposition in the lung as a measure of tissue remodeling following the severe infection-induced injury caused by Nb burrowing through the lung during the acite infection stages. Immunofluorescent staining for Collagen VI was performed on lung sections from vehicle and THC-treated naïve and infected mice (Fig. 8A). Vehicle-treated infected mice showed reduced Collagen VI staining compared with vehicle naïve controls, consistent with infection-associated disruption or remodeling of the lung extracellular matrix^9,16,18^. In contrast, Collagen VI signal was not significantly reduced in THC-treated infected mice compared with THC-treated naïve controls (Fig. 8B). Together, these data suggest that Nb infection is associated with loss of lung Collagen VI in vehicle-treated mice, whereas THC-treated mice retain Collagen VI following infection. This pattern suggests that THC-associated immune reprogramming is accompanied by altered extracellular matrix remodeling during Nb-induced lung inflammation^51^.

**Figure 8.**
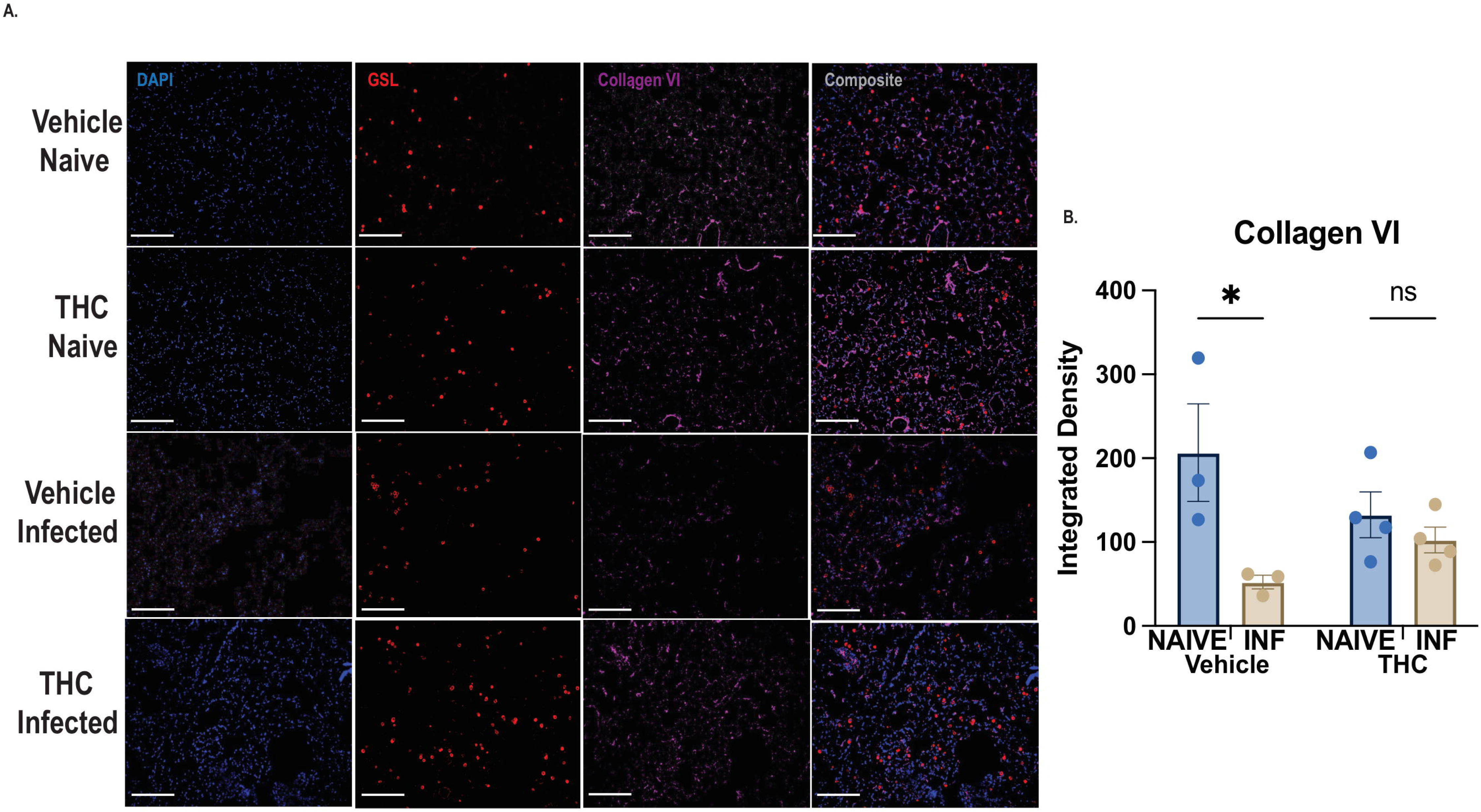
THC treatment is associated with retained lung Collagen VI following Nb infection. Lung extracellular matrix remodeling was assessed in vehicle and THC-treated naïve and Nb-infected mice. (A) Representative immunofluorescence images of lung sections stained for DAPI, GSL I, and Collagen VI. Scale bar = 20 µm. (B) Quantification of Collagen VI integrated density. Data are presented as individual animals with group means ± SEM. Statistical analysis was performed using two-way ANOVA with Tukey’s multiple comparisons test. *p < 0.05; ns, not significant.

## Discussion

The goal of this study was to determine how THC exposure alters immune responses during type 2 inflammation. To achieve a controlled experimental environment, we employed a *Nippostrongylus brasiliensis* infection model and investigated whether THC impacted parasite burden, systemic immune activation, T cell inflammatory function, and lung eosinophil and macrophage-enriched gene expression. We found that THC did not alter infection-associated weight loss, fecal egg burden, or adult worm burden. Instead, THC restrained infection-induced eosinophilia, increased regulatory T cells associated with reduced T-cell derived inflammatory cytokines. Bulk RNA sequencing of lung innate immune cells further showed that THC reduced basal and infection-associated inflammatory programming in CD11c^+^ macrophage-enriched cells and shifted infected lung eosinophils away from migratory and inflammatory activation to an altered metabolic state involving mTOR signaling and oxidative phosphorylation.

An important finding from this study is that THC exposure did not alter *N. brasiliensis* burden. This is notable because human studies have suggested a relationship between cannabis use and helminthiasis. Among Aka foragers of the Congo Basin, urinary THCA levels were negatively associated with parasite infection and reinfection, supporting the possibility that cannabis exposure may influence helminth burden^6^. However, that study evaluated whole cannabis use in a human population, where additional cannabinoids, route of exposure, tobacco co-use, nutrition, behavior, and environmental variables may contribute to parasite outcomes. In our controlled preclinical model, THC exposure alone did not alter egg or worm burden. These findings suggest that purified THC exposure is not sufficient to reproduce the cannabis-associated reduction in parasite burden observed in human populations. Instead, the major effects of THC in this study were immunologic rather than antiparasitic.

Although parasite burden was unchanged, THC exposure altered the host immune response. The reduction in infection-induced circulating eosinophilia is consistent with prior work implicating cannabinoid signaling in eosinophilic inflammation during helminth infection. Inhibition of peripheral cannabinoid receptor subtype 1 during *N. brasiliensis* infection increased airway eosinophilia and worsened pulmonary pathology, suggesting that cannabinoid signaling can restrain eosinophilic lung inflammation^5,10^. Our results indicating reduced circulating eosinophils and decreased inflammatory pathways in lung eosinophils in THC-treated mice are consistent with the broader concept that cannabinoid pathways dampen eosinophil-associated inflammation. RNA sequencing of infected lung eosinophils showed that THC altered eosinophil transcriptional state without reducing lung eosinophil accumulation. THC-treated eosinophils showed increased expression of genes associated with stress adaptation, lipid mediator regulation, and inhibitory receptor signaling, including *Ucp2, Cirbp, Rbm3, Pla2g7, Cyp4f18, Cd300lf,* and *Siglecf*. Increased *Rbm3* is consistent with prior work linking it to restrained type 2 lung inflammation, while *Cirbp* and *Gadd45a* are defined as stress-response genes with context-dependent roles in inflammation^21–23^ Increased *Cyp4f18*, *Pla2g7*, *Cd300lf*, and *Siglecf* is consistent with altered lipid mediator and inhibitory receptor-associated programs in eosinophils, given prior evidence linking CYP4F18 to leukotriene B4 metabolism and CD300/Siglec-family receptors to eosinophil survival, chemotaxis, and effector function^24–28^. In contrast, genes associated with adhesion, migration, and inflammatory stress, including *Macf1, Syne2, Afap1l2, Neat1,* and *Hsph1*, were reduced. These transcriptional changes suggest that THC does not primarily attenuate eosinophilic inflammation by blocking eosinophil entry into the lung.

Instead, THC may alter the activation state of eosinophils that enter inflamed tissue. This distinction is important because eosinophils can contribute to both protective type 2 immunity and tissue pathology depending on their inflammatory context, activation state, and tissue localization^60^.

THC also altered CD11c^+^ lung macrophage-enriched cells before and during infection. Under naïve conditions, THC reduced chemokine, cytokine signaling, and heat shock-associated transcripts in CD11c+ macrophage-enriched cells, while increasing genes associated with stress buffering, lipid handling, and inhibitory signaling. During infection, THC reduced inflammatory, stress-associated, chemotactic, and remodeling-associated genes, including *Tlr4, Fpr1, P2ry13, Hsph1, Hsp90aa1,* and *P4ha1*. Reduced *Tlr4*, *Fpr1*, and *P2ry13* is consistent with dampened pathogen-sensing, formyl peptide receptor, and purinergic inflammatory programs, pathways previously linked to helminth-associated Th2 responses, lung leukocyte recruitment, and macrophage inflammatory regulation^31,38,39,41,45^. Reduced *P4ha1* is also consistent with decreased collagen-processing or ECM-remodeling-associated transcriptional programs^45^. These data indicate that THC changes the baseline state of lung macrophage-enriched cells and alters their response during inflammatory challenge. This is consistent with prior studies showing that THC can regulate myeloid cell biology, including inhibition of macrophage costimulatory activity, alveolar macrophage inflammatory responses, and ROS-associated macrophage differentiation^12–14^. The transcriptional changes observed in macrophage-enriched cells may have implications for tissue remodeling. THC reduced expression of genes associated with motility, inflammatory stress, and fibrotic remodeling, while increasing metabolic and stress-adaptive pathways. Consistent with this transcriptional pattern, vehicle-treated infected mice showed reduced lung Collagen VI staining, whereas THC-treated infected mice retained Collagen VI signal following infection. Collagen VI is a structural extracellular matrix protein, and changes in its deposition may reflect altered tissue remodeling during inflammation. While these data suggest that THC-associated immune reprogramming is accompanied by altered lung matrix remodeling^9,51^.

In CD11c+ macrophage-enriched cells from infected mice, THC-treated samples maintained or increased expression of multiple MHC class II-associated genes, while *Cd80* was reduced. Because CD80-mediated co-stimulation contributes to optimal T cell activation and cytokine production, reduced *Cd80* expression may help explain why CD4^+^ T cells accumulated in the airway but showed reduced inflammatory output after stimulation^49^. The combination of maintained antigen presentation genes, reduced *Cd80*, and reduced TNFα and IFNγ production is consistent with incomplete inflammatory T cell activation or an anergy-like functional state^12,49,50^. These findings are consistent with prior work showing that THC suppresses proliferation, reduces IFNγ production, and alters Th1/Th2 cytokine balance in activated human T cells^11^.

There remain limitations to this study. First, although THC is a partial cannabinoid receptor agonist, we did not directly test whether the observed effects were mediated through CB1, CB2, or cannabinoid receptor-independent pathways^2,12,14^. Future studies using receptor antagonists, conditional deletion mouse models, or cell-specific approaches will be needed to determine receptor dependence. Second, the CD11c^+^ lung population used for RNA sequencing was macrophage enriched but may also contain dendritic cells or other CD11c-expressing populations. Third, although reduced *Cd80* expression and lower TNFα and IFNγ release are consistent with reduced costimulatory activation^11,49,58^, direct lung T cell assays will be required to determine whether THC induces T cell anergy, altered proliferation, or changes in antigen-specific effector function. Lastly, many physiological effects of cannabis are mediated by THC; however, pure THC is rarely consumed on its own, as people typically consume whole cannabis or concentrated cannabis extracts. Future studies should evaluate possible differential effects for THC versus whole cannabis, which contains over 100 phytocannabinoid chemicals^61^(REFERENCE PMID 25315390, Mechoulam et al., 2014).

In conclusion, chronic THC exposure reprograms eosinophils and macrophages toward restrained, oxidative, and stress-adaptive phenotypes during type 2 inflammation associated with reduced inflammatory T cell responses. The immune skewing potential of THC on lung innate immune cells may off therapeutic potential in inflammatory settings where excessive leukocyte activation contributes to tissue pathology^9,16,18^.

## Acknowledgments

We would like to thank the following University of California equipment cores: UC Riverside School of Medicine Research Core for the flow cytometry equipment and biostatistics analysis (somresearch.ucr.edu), UC Riverside High Performance Computing Center for bioinformatics data processing (hpcc.ucr.edu); UC Los Angeles Translational Pathology Core Laboratory for tissue processing. This work was supported by the National Institutes of Health (R01AI153195 to MGN and JJ; R21AI180561, R01AI191470 to MGN) and RAMP Award EDUC5-13636 to KML). The contents of this publication are solely the responsibility of the authors and do not necessarily represent official views of CIRM or other agencies of the State of California.

## Author Contributions

M.G.N, J.J., M.O., N.D.P conceptualized the study and designed the experiments. J.J., M.O., K.M.L., S.M. performed the experiments. J.J., K.M.L. analyzed the data. J.J., N.D.P., M.G.N. interpreted the results. J.J., M.G.N. wrote the manuscript. M.G.N. acquired project funding. All co-authors approved the manuscript.

## Supplemental material

***Supplemental Figure 1*. Flow cytometry gating strategies for whole blood immune cell analysis**. Representative gating strategies are shown for whole blood immune cell populations. (A) Gating strategy for monocytes and eosinophils. (B) Gating strategy for CD4+CD25+ Tregs. (C) Quantification of CD4+ T cells among CD45⁺ cells. Data are presented as mean ± SEM and pooled from 2 independent experiments with n = 4 to 7 mice per group per experiment. Statistical analysis was performed using two-way ANOVA with Tukey’s multiple comparisons test. ***p < 0.001; ****p < 0.0001; ns, not significant.

***Supplemental Figure 2*. Flow cytometry gating strategies for whole lung immune cell analysis**. Representative gating strategies and quantification are shown for whole lung immune cell populations. (A) Whole lung gating strategy for alveolar macrophages and eosinophils (B) quantification of alveolar macrophages among CD45⁺ cells. (C) Quantification of eosinophils among CD45⁺ cells. Data are presented as mean ± SEM and pooled from 2 independent experiments with n = 4 to 7 mice per group per experiment. Statistical analysis was performed using two-way ANOVA with Tukey’s multiple comparisons test. ***p < 0.001; ****p < 0.0001; ns, not significant.

***Supplemental Figure 3*. Flow cytometry gating strategies for BALF immune cell analysis**. Representative gating strategies and quantification are shown for BALF immune cell populations (A) BALF gating strategy for alveolar macrophages, eosinophils, and CD4⁺ T cells. (B) Quantification of alveolar macrophages among CD45⁺ cells. (C) Quantification of eosinophils among CD45⁺ cells. Data are presented as mean ± SEM and pooled from 2 independent experiments with n = 4 to 7 mice per group per experiment. Statistical analysis was performed using two-way ANOVA with Tukey’s multiple comparisons test. ***p < 0.001; ****p < 0.0001; ns, not significant.

---

**Supplemental Table 1.** Whole Lung/Blood Flow Cytometry Panel.

| **Laser** | **Fluorophore** | **Surface** | **Company** | **Cat.** | **Clone** |
| --- | --- | --- | --- | --- | --- |
| 488 nm | FITC | F4/80 | Biolegend | 123108 | BM8 |
|  | PerCP, 7AAD | 7AAD | Biolegend | 420404 | N/A |
|  | PerCP Cy5.5 | CD4 | Biolegend | 100540 | RM4-5 |
|  | PE Cy7 | Ly6C | Invitrogen | 25-5932-82 | HK1.4 |
| 637 nm | APC | CX3CR1 | Biolegend | 149008 | SA011F11 |
|  | AF700 | MHCII | Invitrogen | 50-169-17 | M5/114.15.2 |
|  | APC Cy7 | CD11b | Biolegend | 101226 | M1/70 |
| 405 nm | BV510 | Ly6G | Biolegend | 127627 | 1A8 |
|  | BV570 | CD45 | Biolegend | 103135 | 30-F11 |
|  | BV650 | CD86 | Biolegend | 105036 | GL-1 |
|  | BV711 | CD206 | Biolegend | 141727 | C068C2 |
| 561 nm | PE | CD25 | Biolegend | 113704 | SA203G11 |
|  | PE Dazzle | SiglecF | Biolegend | 155530 | S17007L |
|  | PeCy5 | CD3ε | eBioscience | 15-0031-82 | 145-2C11 |
|  | PE Cy5.5 | CD11c | Invitrogen | 35-0116-42 | 3.9 |

**Supplemental Table 2.** BALF Flow Cytometry Panel.

| **Laser** | **Fluorophore** | **Surface** | **Company** | **Cat.** | **Clone** |
| --- | --- | --- | --- | --- | --- |
| 488 nm | FITC | F4/80 | Biolegend | 123108 | BM8 |
|  | PerCP, 7AAD | 7AAD | Biolegend | 420404 | N/A |
|  | PerCP, Cy5.5 | CD4 | Biolegend | 100540 | RM4-5 |
|  | PerCP eFlour 710 | CD45R | eBioscience | 41-0452-80 | RA3-6B2 |
|  | PE Cy7 | Ly6C | Invitrogen | 25-5932-82 | HK1.4 |
| 637 nm | APC | CD3 | Biolegend | 100236 | 17A2 |
|  | AF700 | MHCII | Invitrogen | 50-169-17 | M5/114.15.2 |
|  | APC Cy7 | CD11b | Biolegend | 101226 | M1/70 |
| 405 nm | BV510 | Ly6G | Biolegend | 127627 | 1A8 |
|  | BV570 | CD45 | Biolegend | 103135 | 30-F11 |
|  | BV650 | CD86 | Biolegend | 105036 | GL-1 |
|  | BV711 | CD206 | Biolegend | 141727 | C068C2 |
| 561 nm | PE Dazzle | Siglec F | Biolegend | 155530 | S17007L |
|  | PE Cy5.5 | CD11c | Invitrogen | 35-0116-42 | 3.9 |

**Supplemental Table 3.** Lung Immunofluorescence Panel.

| **Primary** | **Company** | **Cat.** |
| --- | --- | --- |
| GSL I Isolectin B4, Biotinylated | Vector<br>Laboratories | B-1205-.5 |
| Rabbit anti-mouse COL6A1 PolyAb | Proteintech | 17023-1-AP |
| **Secondary** | **Company** | **Cat.** |
| SAV Cy3 | Invitrogen | SA1010 |
| Goat anti-Rabbit Cy5 | KPL | 072-02-15-06 |

