## Supplementary figures and images for "Δ^9^-Tetrahydrocannabinol exposure shifts eosinophil and macrophage transcriptional programs towards an anti-inflammatory phenotype in helminth infection"

### Supplemental Figure 1

Sup. Fig. 1.

**A.** PBMC - Eosinophils and Monocyte Gating Strategy

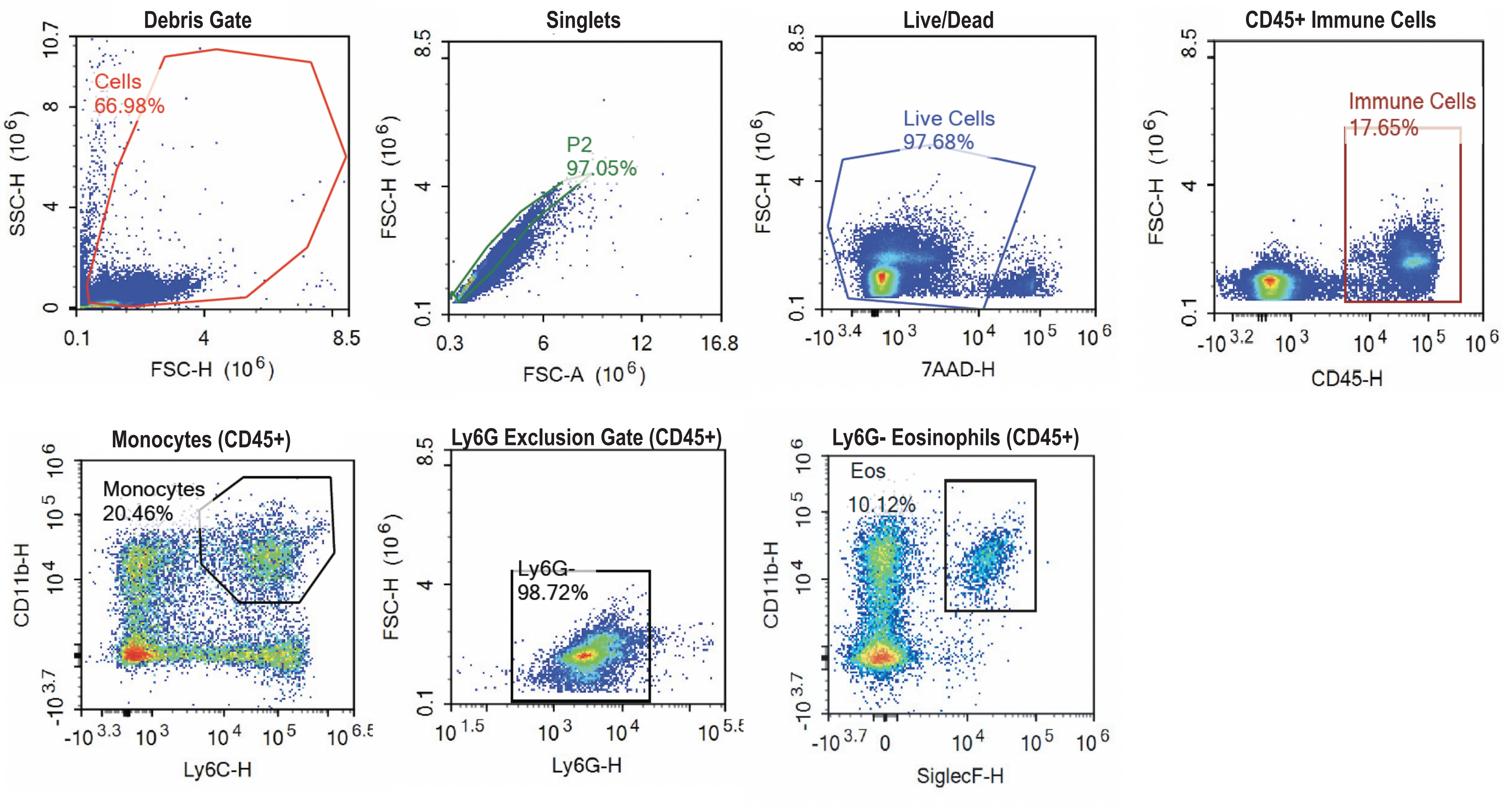


**B.** PBMC - T Cell Gating Strategy

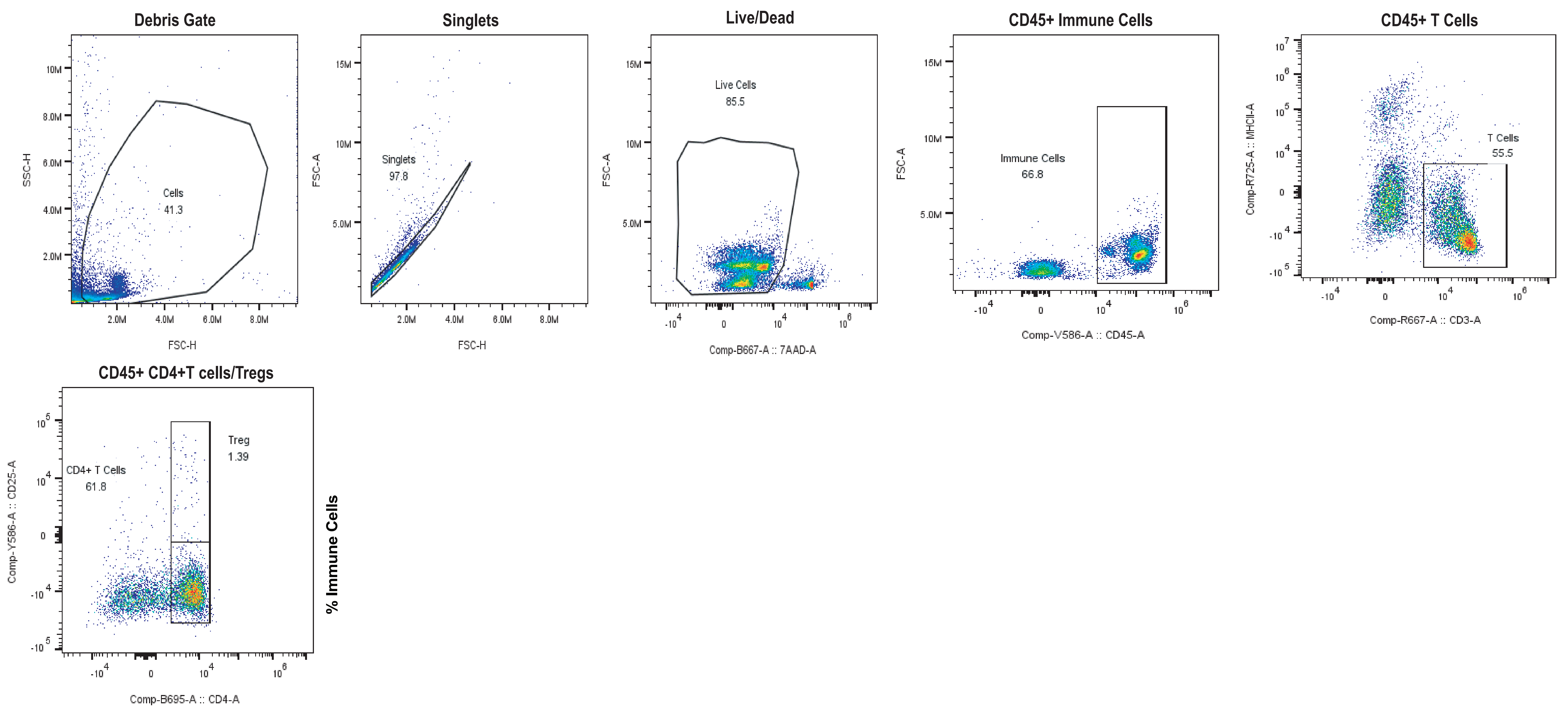


**C.**

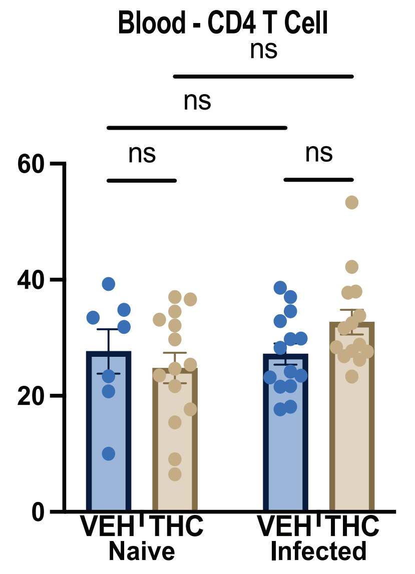

### Supplemental Figure 2

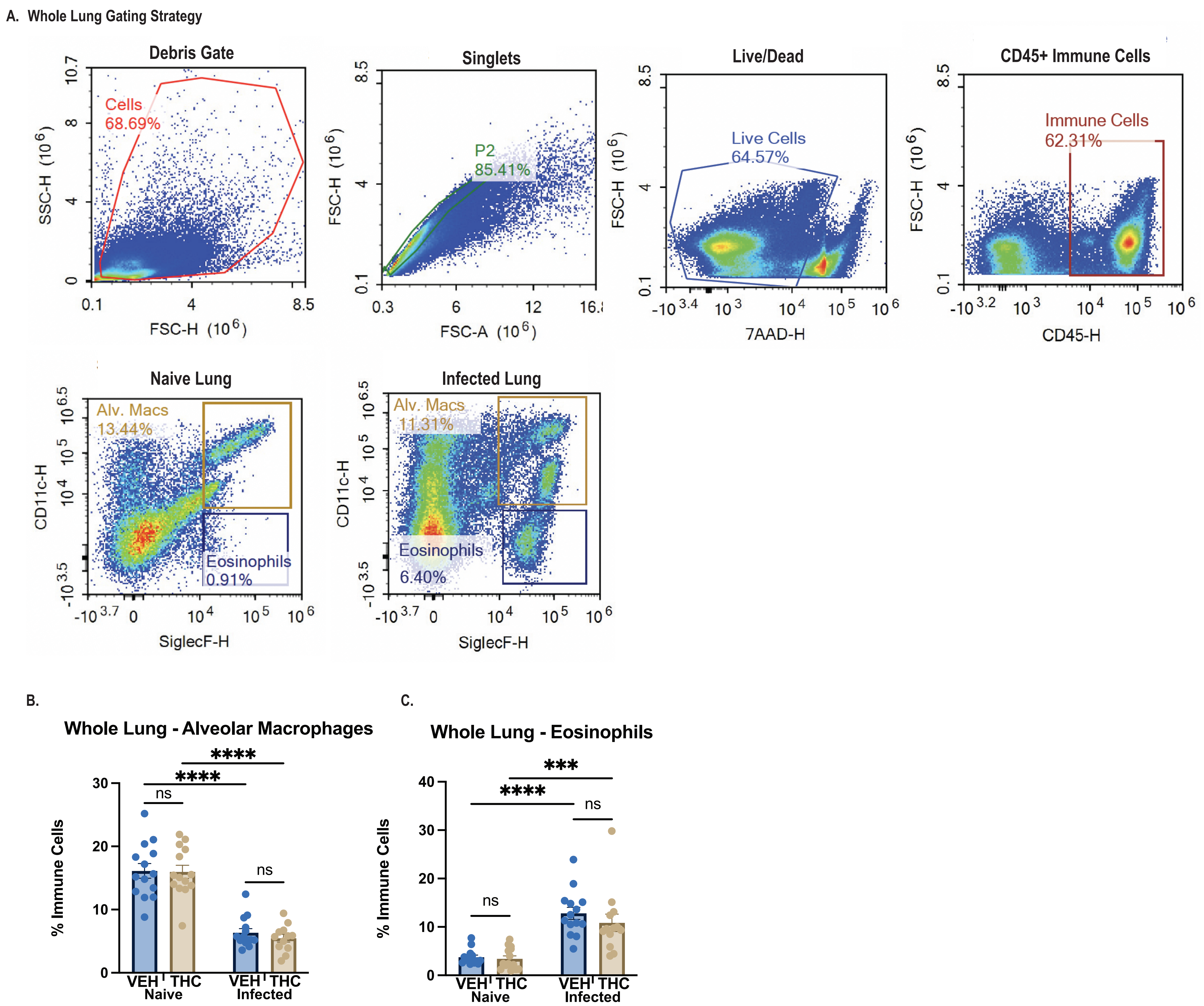

### Supplemental FIgure 3

Sup. Fig. 3.

A. BALF Flow Cytometry Gating Strategy

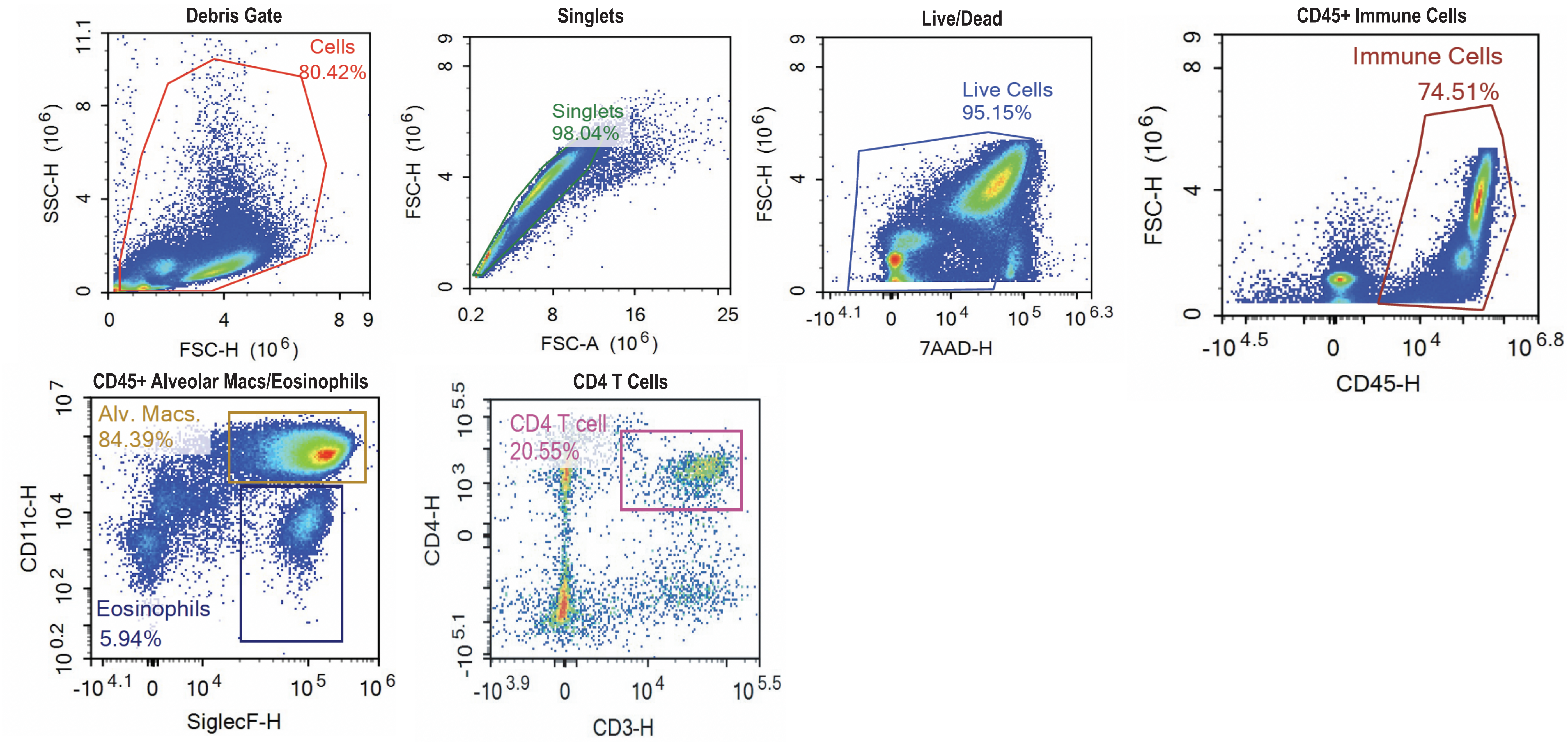


B.

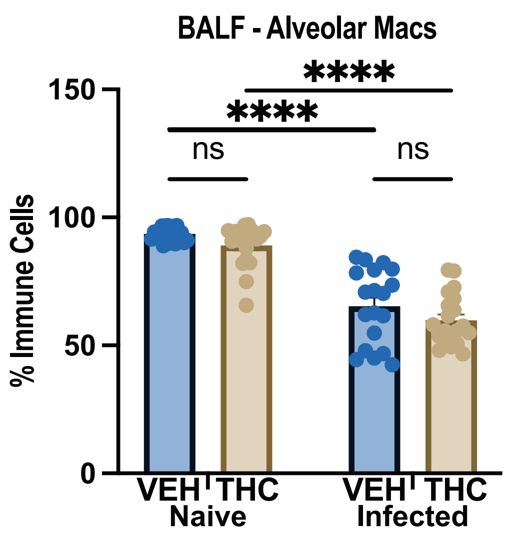


C.

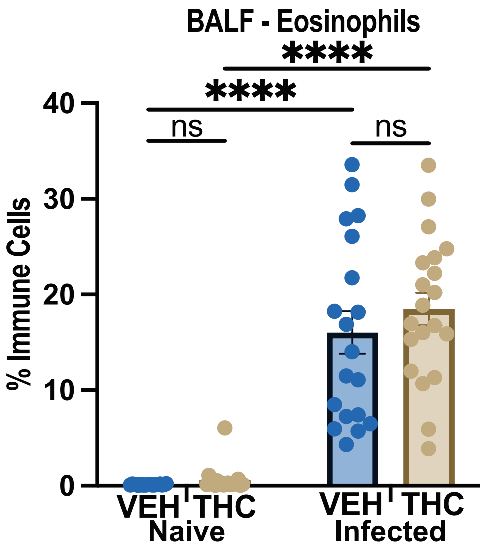
